# Hybrid Exb/Mot stators require substitutions distant from the chimeric pore to power flagellar rotation

**DOI:** 10.1101/2024.03.12.584617

**Authors:** Pietro Ridone, Matthew A. B. Baker

## Abstract

Powered by ion transport across the cell membrane, conserved ion-powered rotary motors (IRMs) drive bacterial motility by generating torque on the rotor of the bacterial flagellar motor. Homologous heteroheptameric IRMs have been structurally characterized in ion channels such as Tol/Ton/Exb/Gld, and most recently in phage defense systems such as Zor. Functional stator complexes synthesized from chimeras of PomB/MotB (PotB) have been used to study flagellar rotation at low ion-motive force achieved via reduced external sodium concentration. The function of such chimeras is highly sensitive to the location of the fusion site, and these hybrid proteins have thus far been arbitrarily designed. To date, no chimeras have been constructed using interchange of components from Tol/Ton/Exb/Gld and other ion powered motors with more distant homology.

Here we synthesised chimeras of MotAB, PomAPotB and ExbBD to assess their capacity for cross-compatibility. We generated motile strains powered by stator complexes with B-subunit chimeras. This motility was further optimised by directed evolution. Whole genome sequencing of these strains revealed that motility-enhancing residue changes occurred in the A-subunit and at the peptidoglycan binding domain of the B-unit, which could improve motility. Overall, our work highlights the complexity of stator architecture and identifies the challenges associated with rational design of chimeric IRMs.

**Importance:** Ion-powered rotary motors (IRMs) underpin the rotation of one of nature’s oldest wheels, the flagellar motor. Recent structures have shown that this complex drives ion conduction and even phage defence and thus appears to be a fundamental molecular module with diverse biological utility where electrical energy can be coupled to rotational force to execute work.

Here, we attempted to rationally design chimeric IRMs to explore the cross-compatibility of these ancient motors. We succeeded in making one working chimera of a flagellar motor and a non-flagellar transport system protein. This had only a short hybrid stretch in the ion-conducting channel, and function was subsequently significantly improved through additional substitutions at sites distant from this hybrid pore region. Our goal was to test the cross-compatibility of these homologous systems and highlight challenges arising when engineering new rotary motors from this fundamental nanomotor.

## Introduction

Many bacteria are propelled by the rotation of flagella to navigate to better environments. The rotation of these flagella is powered by ion-powered rotary ion-pumps known as stators(1–3). Recent structural characterization of the stator elements that are ion conductors revealed that these are heteroheptamers (MotA_5_MotB_2_(1, 2)), and this has now been observed in sodium powered stators (PomA_5_PomB_2_(3)), in ion pumps (GldM_5_GldN_2_(4), ExbB_5_ExbD_2_(5–7)) and most recently in phage defense systems (ZorA_5_ZorB_2_(8)). These ion-powered rotary machines (IRMs)(9) have diverse functions but high structural conservation implying a common origin. Thus, IRMs represent a key molecular complex with an origin at least as ancient as flagellar motility(10, 11). As such there is significant interest in understanding what the structure and function of a potential common ancestor might be.

The flagellar motor itself is highly modular (Fig.1). This organization is routinely described as the filament, hook and basal body(12), an oversimplification which does not even include the IRMs as a separate module. We consider submodules of the motor to be identifiable domains that have a specific function: in the case of the stator-complex, this is commonly defined for the central B-unit as the transmembrane (TM) domain, the linker, and the peptidoglycan-binding domain (PGB, in gram negative species such as *E. coli*). As more structural information becomes available, it is also possible to clearly identify pore-facing, rotor-interacting, and lipid-facing domains in the pentameric A-unit(13–15).

**Figure 1.**
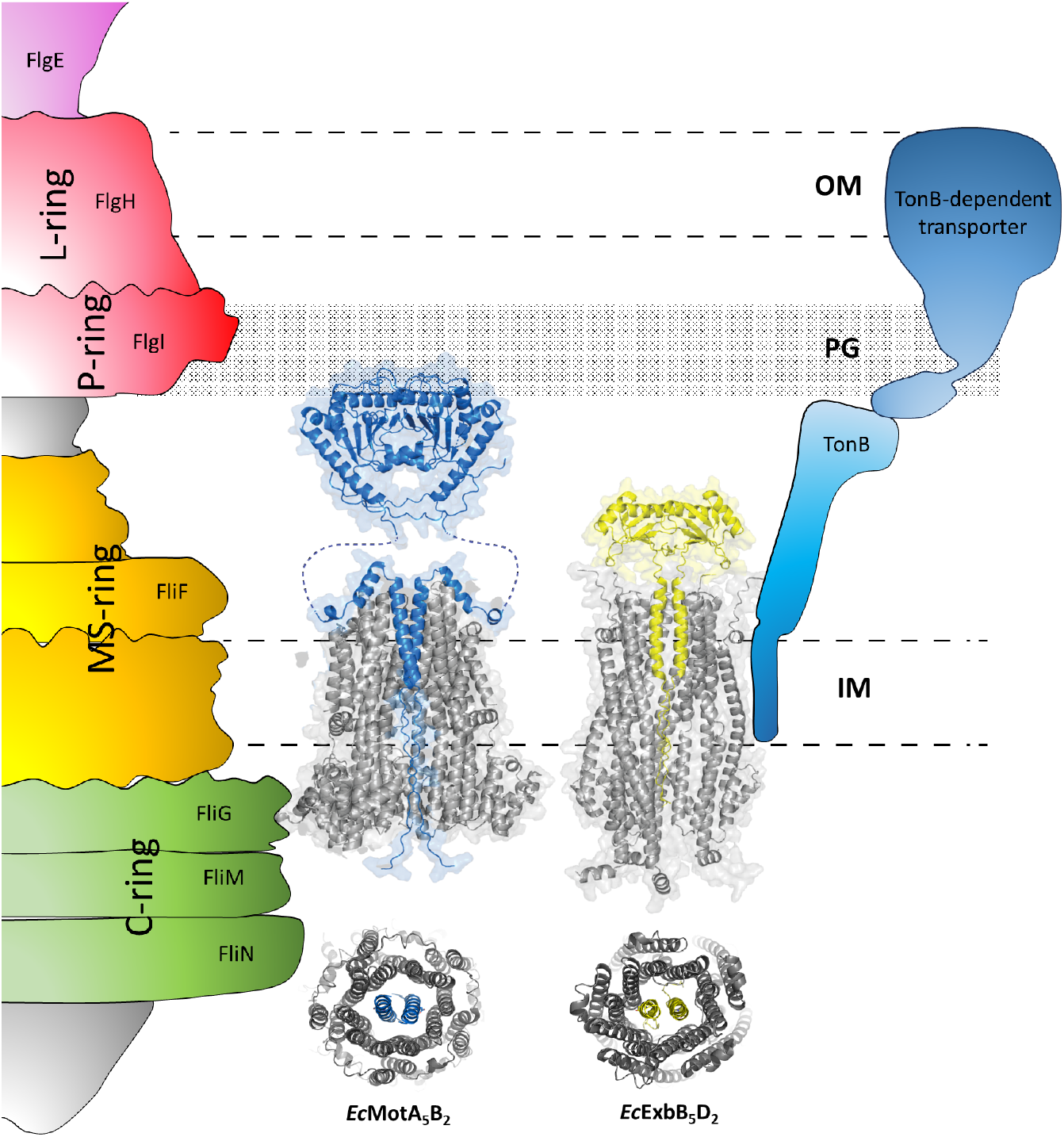
Overview of *E. coli* MotAB and ExbBD complexes. The Bacterial Flagellar Motor and the TonB-dependent siderophore uptake system of *E*.*coli* are powered, respectively, by two homologous heptameric systems, the MotA MotB and the ExbB ExbD complexes. These two transmembrane systems share their general architecture, subunit stoichiometry and functional features, as both depend on interaction with the peptidoglycan (PG) layer and the presence of an ion-motive force (IMF) to operate. The MotAB complex interacts with FliG at the cytoplasmic interface of the basal body of the BFM (left) while the ExbBD complex is thought to interact with the TM domain of TonB at the plasma membrane.

Chimeric work in the stator complex is not new: the first Mot/Pom chimeras were designed in the 1990s with the target of generating a sodium-powered chimera that was functional in *E. coli* (16, 17). Mutagenesis and protein engineering work exploring the role of the plug, linker and peptidoglycan-binding domains in light of new structures continues(18–23) as well as efforts to reconstruct an ancestral stator to explore origins and divergence(14, 24). However, to our knowledge no attempt has been made to combine homologous A_5_B_2_ pumps coupled to non-flagellar function with existing stator components.

In this work, we seek to explore the possibility of engineering functional stator chimeras. We explore compatibility of submodules of IRMs and test these for their capacity to restore motility in stator-deleted bacterial strains. We show the capacity of the overall flagellar complex to rescue motility through epistatic substitutions that affect residues far from chimeric interfaces, revealing the complex integration of submodules across the complex. In doing so, we explore the limits of function as well as the potential for customizing new functions and determining essential protein motifs which differentiate structure and function across the IRM classes.

## Results

We designed chimeras of *motA* and *exbB* (the A-subunit) and *motB exbD* (the B-subunit) by performing amino acid alignments between homologs and by manual inspection of their 3D structures to select the residues connecting the chimeric portions of our constructs. We selected 4 A-subunit and 11 B-subunit constructs to clone into a plasmid vector (pBAD33) for *in vivo* testing in a stator-deleted strain (*E. coli* Δ*motAB*) (SI Fig. 1A, 1B).

Each construct was maintained in culture on a soft agar swim plate for up to 2 weeks to allow for the isolation and propagation of up-motile populations. Out of all 14 tested constructs we isolated one *potB / exbD* variant (PotD_3.3_) able to restore motility in presence of *pomA* (Fig. 2). The plasmid carried by this motile population was extracted and both stator genes were sequenced to check for additional mutations contributing to the phenotype. A missense mutation (*CAA* to *CCA*) causing the Q104P substitution was found in the plasmid-encoded chimeric construct *potD*_*3*.*3*_. (SI Fig. 2). This stator B-subunit construct (*potD*_*3*.*3*_ *Q104P*) was renamed “PotDα”.

**Figure 2.**
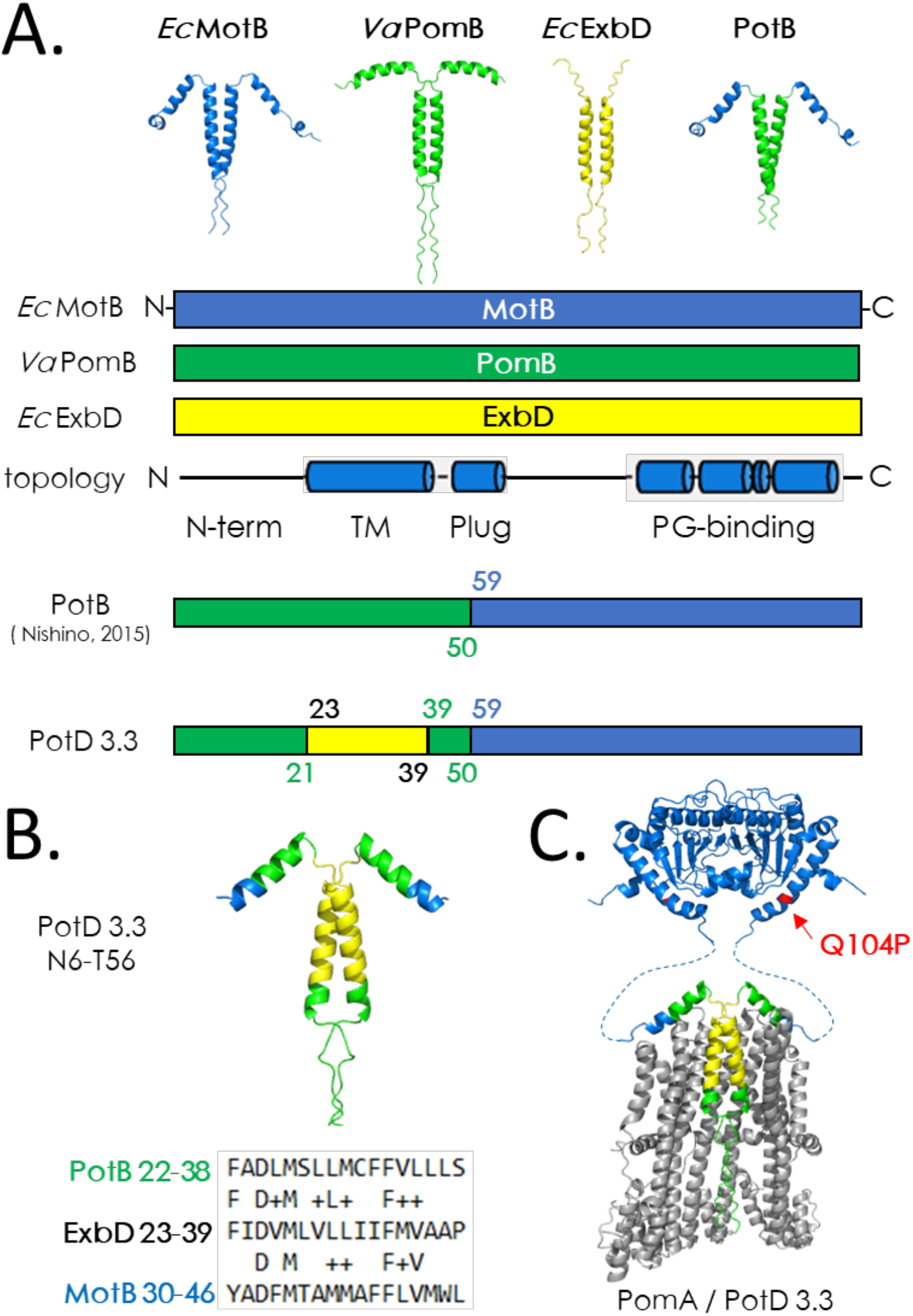
Isolation of an ExbD/stator chimera able to drive motility . A) Representative Alphafold2 models of the TM and plug portions of *Ec* MotB (*Escherichia coli*, blue), *Va* PomB (*Vibrio alginolyticus*, green), *Ec* ExbD (yellow) and chimera PotB. Schematic diagrams of each B subunit, their general topology (with alpha helices depicted as cylinder and sections lacking a secondary structure as connnecting lines) and junction site annotations for the previously reported PotB chimera and candidate PotB/ExbD chimera PotD 3.3. Coloured sections of the diagrams indicate which original protein the fragment belongs to in the chimeric construct. Numbers flanking each junction site indicate the residues belonging to each portion of the chimera and are color-coded according to the scheme above. B) Representative model of the TM and plug portion (N6-T56) of PotD 3.3, coloured according to the scheme in A. A sequence similarity alignment is shown to highlight the degree of conservation between the TM domains of ExbD (residues 23-39) and PotB (22-38), and ExbD and MotB (residues 30-46). Letter intercalating the alignment indicate identity of homologous residues while the ‘+’ sign indicates similarity. C). The position of suppressor mutation Q104P in PotD 3.3 is mapped (red) on a composite heptameric (A_5_B_2_ ) model of a PomA (grey) /PotD 3.3 (coloured as in B) stator.

To verify that Q104P was the substitution that rescued motility, we deleted *pomA* from the mutant plasmid and tested the function of PotDα in combination with various A-subunits. PotDα was tested in presence of *motA* and all chimeric A-subunit constructs to check whether together they could form a functional stator (Fig. 3A, 3B). Interestingly, none of our strains expressing *potDα* / A-subunit combinations produced swim rings within 24 hr (SI Fig. 3A) whereas cells containing control combinations (*motAmotB, motApotB* and *pomApotB*) all formed swim rings on plates and produced observable swimming cells (Fig. 3C) powered by proton (*motA*) or sodium (*pomA*). Thus, none of our A-subunit chimeras functioned in presence of any B subunit partner, whether chimeric or wild-type. Notably, bacteria co-expressing the *mobB*.*v1 potDα* combination showed a slower growth phenotype when compared to *motAmotB* control (SI Fig.3B), suggesting that the stator construct might leak ions although being unable to generate torque.

**Figure 3.**
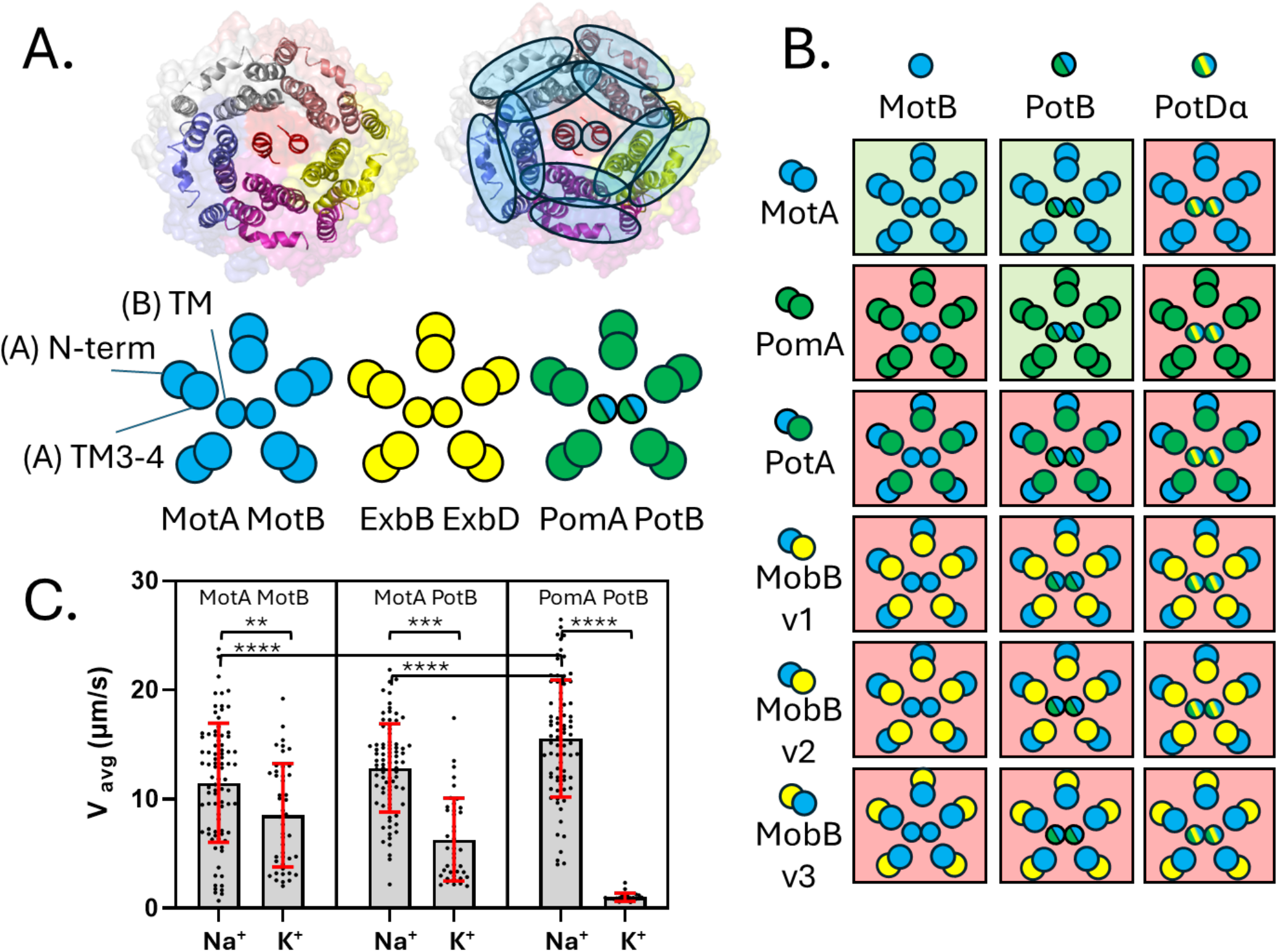
Functional characterization of A-B chimeric interfaces. (A) Section of the stator complex A-B interface viewed from the periplasmic side. Monomers of MotA, shown in different colors, and blue ovals highlight the pore-facing (interior) and lipid-facing (exterior) residues of the complex. Circular diagrams are used to illustrate the intended arrangement of each protein subdomain in the chimeras tested for their ability to rescue motility. (B) Summary of all motility tests performed in stator-deleted *E. coli* co-transformed with plasmids bearing an A-subunit construct (rows) and a B-subunit construct (columns). The background color of each box indicates a motile (green) or non-motile (red) phenotype. Each A-B combination is schematically represented using the symbology shown in A. (C) Single-cell free-swimming cell velocity measurements for motile strains expressing the stator complexes highlighted in green in (B), expressed from plasmid in *E. coli* Δ*motAB*. Cell velocity was measured in sodium-based (85mM NaCl) and potassium-based (85mM KCl) LB medium. (N = 84 (MotA MotB, Na^+^), 45 (MotA MotB, K^+^), 85 (MotA PotB, Na^+^), 41 (MotA PotB, K^+^) and 77 (PomA PotB, Na^+^), 24 (MotA PotB, K^+^) cells). Error bars indicate the standard deviation of the mean. Statistical comparison was performed using Student’s T test, ** P < 0.01; *** P <0.001; **** P < 0.0001.

Because the Q104P substitution alone was unable to rescue motility, we screened for improved swimmers by directed evolution to isolate additional mutations in stator genes that might have contributed to phenotypic rescue. We subjected our motile PotDα isolate and all non-motile, B-subunit chimeras to repeated passaging on fresh swim plates at 3-4 days intervals. Initially, swim rings of the *pomApotDα* strain spread noticeably slower than the *pomApotB* strain but after six passages motility had improved considerably. Visible swim rings of *pomApotDα* formed within a 24-hr period (SI Fig. 4A) compared to the multiple days required for the original flare (SI Fig. 2). The *potDα* variant was motile not only in the presence of both *motA* and *pomA* but also when only *pomA* was present (tested in Δ*motB E. coli* RP3087, *motB* on genome, *pomA* on plasmid; or *E. coli* Δ*motAB*, only *pomA* from plasmid). Plasmid sequencing of this variant revealed an additional mutation in the plasmid-encoded *pomA* gene (causing the substitution A86V). All other B-subunit constructs, which did not support motility, showed no rescue of motility after 6 passages.

We used genome-based directed evolution of motility(25) to attempt to further improve motility in our chimeric variant. We replaced the native *motAB* locus of *E. coli* RP437 with the *pomApotDα* construct cloned from the plasmid (SI Fig.4A, 4B). As observed in directed evolution experiments of strains expressing the chimeras from a plasmid vector, flares developed over multiple days, and motility improved after several passages (SI Fig. 4B.) A control genomic edit using the original *potD*_*3*.*3*_ construct (*E. coli* “PotD”, SI Fig.5A, 5B), without additional mutations, did not adapt to rescue motility (SI Fig. 5C). Whole-genome sequencing of members of our PotDα evolved lineage revealed a new missense mutation in the chromosomal *pomA* gene (substitution H116Y) and additional mutations in the *flgM* and *icd* genes (SI Fig. 4B, SI Table 2).

To repeat the effect of these stator mutations, we introduced them into the plasmid-encoded *pomApotDα* constructs via site-directed mutagenesis and measured their effect on motility in our stator-deleted strain (*E. coli* Δ*motAB*). Growth profiles of all our evolved and engineered strains showed no effect on growth when *potDα* was expressed from plasmids, while growth was inhibited when the construct was expressed from the edited locus on the chromosome (SI Fig. 6A). At the same time, we tested the effect on swim rings in the presence and absence of sodium to determine the ion-selectivity of the chimeric stators (Fig. 4A, SI Fig. 6B). Our strains expressing chimeric stators appeared to require Na^+^ for effective motility and did not produce swim rings on K^+^ swim plates. Motility was inhibited in presence of the Na^+^-powered stator blocker Phenamil (SI Fig. 6B). Our plasmid-based validation showed that the A86V/H116Y residue changes in PomA improved the motility phenotype when co-expressed with *potDα* (Fig. 4B). Although the Q104P change in PotD_3.3_ was located at the N-terminal end of the PG-binding domain, both *pomA* mutations affected residues on the cytoplasmic side of the stator near the surface that interacts with the rotor (FliG, YcgR). Thus, mutations that improve motility can affect residues located far from each other in the folded protein structure, indicating that they improve different aspects of stator function.

**Figure 4.**
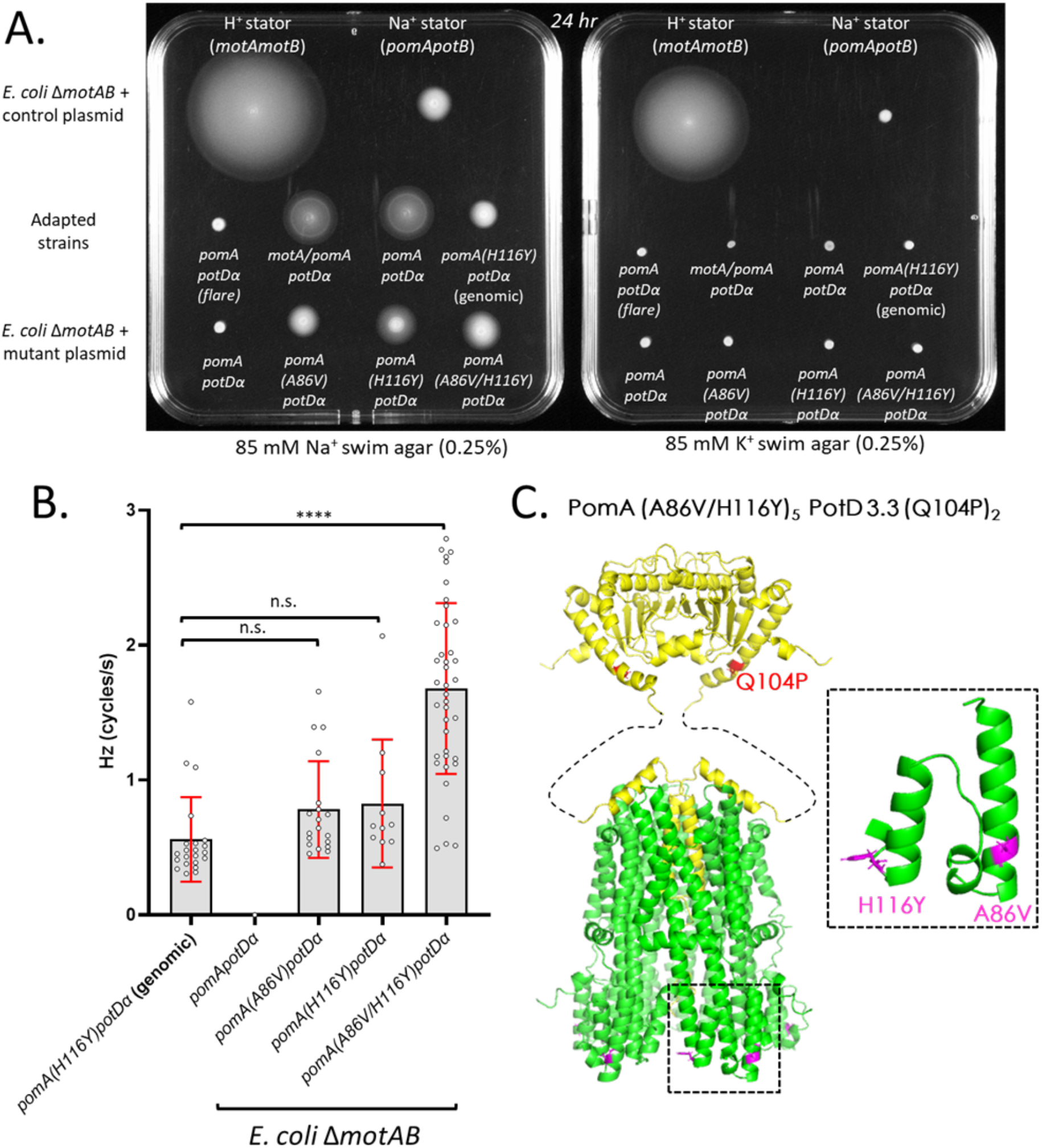
Functional chimeras require sodium for function. (A) Comparison of motile phenotypes of *E. coli* strains. Stator-deleted Δ*motAB E. coli* harbouring a proton-powered (MotAMotB) or sodium-powered (PomA/PotB) stator genes are shown as motility controls at the top of each square petri dish, each supplemented with 85 mM NaCl (left) or 85 mM KCl (right). (B) Rotation speed of single tethered cells expressing PotDα and PomA variants engineered with other candidate motility-enhancing mutations detected in directed evolution experiments. The first bar on the left (PomA(H116Y)PotDα genomic) indicated the construct being expressed from the chromosomal *motAB* locus, while all other constructs were expressed from a plasmid in a *E. coli* Δ*motAB* background (N = 22, 0, 19, 11, 39 cells, respectively; **** P < 0.0001; n.s.: no significant difference). (C) Alphafold2 model of the chimeric stator PomA(A86V/H116Y)_5_PotDα_2_ with A-subunits in green and B-subunits in yellow. The position of each mutation is highlighted and labelled in red or purple on the structure and in the inset.

## Discussion

We engineered 11 chimeras of the B subunit of the hybrid PomAPotB stator with the ExbD component of the ExbBD stator. These hybrids had varying extents of ExbD, ranging from 5.3% to 16.95 %. Of these, the PotD_3.3_ construct with the smallest contribution from ExbD was able to generate torque and rescue motility following subsequent adaptation with directed evolution. None of our A-subunit chimeras were functional or rescued by directed evolution. However, some mutations in *pomA* improved motility when expressed with *potD*_3.3_. These mutations affected regions of PomA that are far from the ion pore but proximal to the stator-rotor interface.

### Fusion-sites are critical in the B-unit

It is challenging to determine why some chimeras are functional where others are not, especially considering that some B-unit chimeras differed only very slightly. Chimeras that differed by only 4 residues (PotD_3.0_), in non-conserved regions adjacent to the pore were unable to rescue motility. This is indicative of the general challenge of rational design and the need to select fusion sites carefully and attempt multiple variants simultaneously to improve throughput. Single amino acid replacements within the TM domain of *Ec*MotB are likely to disrupt motility even when they are conservative(26).

### Preservation of the pore proximal A-B interface does not guarantee function

To date, no functional A-unit chimeras have been made, either from Mot/Pom complexes or across the ion-powered rotary motors more widely. Here, we matched A-B interfaces across the TM region of *Ec*MotB (residues I28-W45) and TM3/4 in *Ec*MotA (residues N139-N250) (SI Fig. 7A, 7B). Despite this, none were functional in driving torque-generation and motility. One combination of chimeric variants aimed at reconstituting a complete ExbBD-like pore (*mobB*.*v1 potDα*) caused growth-delaying effects when expressed, suggesting that it may be able to conduct ions without coupling this current to torque generation (SI Fig. 7C, 7D). Construct MobB v2, which contains the C-terminal domain of ExbB (SI Fig. 1), did not show any growth defects when it was expressed, suggesting that the C-terminal residues of MotA (residues 247-295 of *Ec* MotA) are required for ion conduction(3, 27).

The pore interface of PomA is compatible with both PotB and PotDα, but not with MotB, PotD 4.3 (SI Fig. 1) or a PotB variant expressing the TM segment of MotB (PotB-H^+^, SI Fig. 8), suggesting that the PomB portion of the chimera is crucial to allow PotDα to function in a sodium-dependent fashion. The MotAB stator thus appears to be less likely to allow functional chimeras with ExbD compared with the Pom stator. When testing the function of our chimeric variants we did not investigate whether the inability of some constructs to rescue motility was due to poor expression via Western blot or RT-PCR.

Despite this, our results indicate that regions outside the pore are important in improving engineered pores. However, when the lipid and rotor interfaces were preserved by using a single A-subunit with known functional domains in these locations (*mobB*.*v1, mobB*.*v2* and *potA*, Fig. 2A), these chimeras were still non motile.

### Allosteric substitutions further improve motility in partially functional stators

Motility was rescued in *potD*_*3*.*3*_ via an amino acid substitution at a residue (Q104) located in a different protein domain than the chimeric TM pore domain. The substitution introduced a proline on the first alpha helix of the large PGB domain. This substitution might contribute to the stability of the binding to the peptidoglycan layer and the mechanosensitivity of the chimeric stator. A previously reported substitution (R109P) found in a linker-deleted PotB variant rescued the motility of the host(28), emphasizing that the PGB-linker connection is probably structured.

Subsequent mutations affecting sites far from the hybrid region improved motility and caused residue substitutions near a known FliG interaction site of the A-subunit (PomA) at A86 and H116. Regions far from the pore not only affect function but also repair a hybrid pore to enable function. Substitutions affecting sites far from the pore have been shown to affect phenamil binding and resistance(29), putatively allowing a defective stator to remain engaged with the rotor for longer. It is possible that stabilizing the PG-binding of the stator complex would enable motility driven by a partially functional stator. Such allosteric interactions undermine attempts to rationally design proteins through edits at only a single functional site. Methods to examine allosteric networks across proteins can help these attempts (30–32). Mutations in class III flagellar gene repressor FlgM (anti-FliA, σ^28^ factor) have been shown to impact motility(33), but whether the FlgM S7Y substitution affected the efficiency of FlgM secretion remains to be determined. The Ser7 residue in FlgM is distant from the interaction interface between FliA and FlgM (SI Fig. 9) and might not affect the ability of FlgM to bind to FliA.

### Limitations of testing chimeric constructs in gram-negative *E. coli*

We sought to design chimeras that preserved ion conduction, torque generation and mechanosensitivity and approached this via engineering conservative chimeras centered around the pore. The stator complex has multiple functions, executed by specific domains of the complex that are either pore-facing (TM3-4 residues N139-N250 in *Ec*MotA), rotor-facing (residues L81-M138) or PG-binding (residues N102-L288 in *Ec*MotB)(23). Not only must the stator complex sense force to activate and transduce chemical potential into torque, but it must also interact with the rotor to couple torque to filament rotation. The rotor itself is a large molecular complex with multiple functions that include the conformational rearrangements necessary to change the direction of rotation as well as the conformational changes necessary to generate torque to drive rotation.

All chimeras in our work were only tested in an *E. coli* chassis. Motors in different species have different protein and membrane composition, different sizes, as well as different stator dynamics(34–36). Thus, it is possible our stators are functional but require rotor and membrane environments that are better matched to their chimeric pores. Single-molecule *in vitro* bilayer measurements combined with electrophysiology(37, 38) could provide a platform to assay the role that lipid composition and membrane environment have in affecting the diffusion and ligand-binding of the IRM. This could elucidate which submodules of the motor retain partial functionality, whereas in our *in vivo* motility assays we are only able to test overall function as presence or absence of motility.

### Directed and experimental evolution improves function rapidly

As in our previous work, during iterative swim plating we observed mutants that upregulate filament expression(25, 39). Our plasmid-based and genome-directed evolution generated convergent results. Substitutions affecting residues far from the pore rescued motility within days. These screens allow us to explore epistatic interactions that rescue motility, such as non-pore residues improving function in environments depleted of ions for conduction. However, a simple swim plate does confound growth, motility and chemotaxis, so it does not select only for mutations that alter the capacity for the stator complex to generate torque(25). Nevertheless, we do see substitutions that repair edits and improve performance on swim plates. These effects can be correlated with other measurements, such as tethered cell assays(40–42).

More specific screens that directly target motility(43) or cell energetics could be applied to select for mutations that affect the function of specific submodules of the stator complex. Likewise, *in vitro* methods using purified membrane protein complexes could facilitate determination of the *in situ* structure of the stator within the membrane(44), or used to measure current flow and other biophysical parameters(5). Fluorescently labelled stators could be visualized *in vitro* to detect conformational changes at the single-molecule level(45, 46). Improved preparation of combinatoric libraries using newer approaches in synthetic biology could improve throughput(47–50). We utilized medium throughput biological methods that are still able to select one-in-a-million up-motile variants(29). Future high-throughput screens will require automated colony passaging by liquid-handling or colony-picking robots.

### Implications for the evolution and common ancestry of IRMs

Our experimental results might reflect a closer evolutionary relationship between the Exb and Pom systems than Exb and Mot. The Exb system is nevertheless largely incompatible with the BFM stator systems given how many of our chimeras failed to confer motility. An in-depth phylogenetic and structural assessment of the IRM protein family will be required to confirm this hypothesis.

## Conclusions

Our work shows that even with as few as 12 amino acid substitutions, cut-site location and chimeric design is challenging and requires high-throughput approaches to succeed. Recent structural characterization of structural homologues of the stator complex, including Zorya, has shown the diversity of biological applications where IRMs can transduce energy from ion gradients to mechanical work. Knowing the structure of the common ancestor of these IRMs would shed light on the origin of the stator complex, as well as other complexes, and this may be a key innovation of function in early life.

Construction of chimeras exchanging components between extant homologues does not necessarily inform the structure and function of this common ancestor, but it does allow us to examine what are the core elements of these complexes, as well as understand which modules are interchangeable and which are not. This can inform the order of events under which the stator complex arose and diversified and inform parsimony approaches to loss and gain of function in subsequent adaptation. Further studies with new methods in statistical and structural phylogenetics are required to target ancestors of the stator complex for characterization both structurally and biophysically.

## Materials and Methods

### *E. coli* strains, plasmids, and culture media

Synthetic stator chimeras were tested in stator-deleted derivative strains of *E. coli* RP437, either *E. coli* Δ*motB* (RP3087(51) or *E. coli* Δ*motAB* (SI Fig. 5). For genomic edits, *E. coli* RP437 was used as the parent strain(52) to with the native *motAB* locus edited using No-SCAR methods, as previously (40). The pSHU1234 (pBAD33 backbone, Cm+) plasmid encoding *pomA* and *potB* (29) was used as the vector to clone and express all chimeric constructs.

Chimeric constructs were designed *in silico* and synthesized by Integrated DNA Technologies (IDT) as gene fragments (gBlocks). Chimeras were cloned in plasmid pSHU1234 using restriction digestion with NdeI and PstI (New England Biolabs). Plasmids carrying chimeric constructs were used as templates to generate the double-stranded donor DNA used to replace the *motA* and *motB* gene on the RP437 chromosome with *pomA* and *potD* variants, respectively (see Scar-less editing).

Point mutations in plasmids were generated using the QuikChange technique. Bicistronic plasmids expressing both A and B stator subunits were converted to single-gene plasmids expressing only the B-subunit via inverse deletion PCR. Single-construct plasmids expressing A-subunits were constructed via restriction digestion (NdeI and PstI, New England Biolabs) and ligation using pBAD24 (Amp+) as a plasmid backbone. The PotB TM replacement with the MotB TM domain (PotB-H^+^ construct) was generated by inverse deletion PCR followed by self-ligation of the linearized PCR product. Cloned constructs were confirmed by Sanger sequencing (Ramaciotti Centre for Genomics, Kensington). Gene expression from all plasmids used in this work was induced using 1 mM L-(+)-Arabinose (A3256, Sigma-Aldrich) in liquid and solid media.

Liquid cell culturing was done using LB broth (85 mM NaCl or 85 mM KCl, 0.5% yeast extract (70161, Sigma-Aldrich), and 1% Bacto tryptone). Cells were cultured on agar plates composed of LB broth and 1% Bacto agar (BD Biosciences, USA). Swim plate cultures were performed on the same LB broth-based substrates adjusted for agar content (0.25% Bacto agar). Inhibition of Na+-dependent motility was performed using 100 μM phenamil methanesulfonate (P203, Sigma-Aldrich).

### Scar-less editing of *E. coli* with Cas9-assisted recombineering

This procedure was adapted from the no-SCAR method(53). The target strain to be edited (E. coli RP437) was sequentially transformed first with the pKD-sgRNA-3′*motA* (Sm+) plasmid, encoding a single guide RNA (sgRNA) sequence directed at the 3′ end of *motA*, and then with the pCas9cr4 (Cm+) plasmid to yield a parent strain harboring both plasmids. The overlap extension PCR technique(54) was used to assemble linear double-stranded DNA (dsDNA) molecules using two (knock-out) or three (knock-in) starting dsDNA fragments. The resulting donor DNA was electroporated in the plasmid-bearing host, and the successfully edited clones were selected via colony PCR and Sanger sequencing of the *motAB* locus. A list of oligonucleotide primers and synthetic gene fragments (gBlocks) used in this work is provided in Supplementary Table 1.

### Free-swimming assay preparation and analysis

Motile cells were grown overnight in tryptone buffer (TB: tryptone, 85 mM NaCl) at 30°C to OD600 of ∼0.5, diluted 100x in TB and loaded for imaging in a tunnel slide. A custom LabVIEW software was used as previously(24, 29, 40) reported to measure the speed of swimming cells. Time-lapse videos were collected using a camera (Chameleon3 CM3, Point Grey Research) recording 20-s-long videos at 20 frames/s. Visualization of the data was performed using GraphPad Prism 8.

### Sequencing and genomics

WGS of *E. coli* strains was performed using a MiSeq 2 × 150–base pair (bp) chip on an Illumina sequencing platform. Sequencing was carried out at the Ramaciotti Centre for Genomics, Kensington and delivered as demultiplexed fastQ files (quality control: >80% bases higher than Q30 at 2 × 150 bp). The SNP calling and analysis were performed using Snippy(55–57). The short reads from sequencing were aligned to the MG1655 reference *E. coli* genome (GenBank: U00096.2) and an *E. coli* RP437 (GenBank: CP083410.1) as previously (25). A summary of all single nucleotide polymorphisms (SNPs) detected in this study is provided in Supplementary Table 2.

### Structural modelling

Sequence alignments were aligned using MAFFT. Models of stator monomers and heptameric complexes was performed using AlphaFold2 on Galaxy.au, using the Australian Alphafold Service (AAS). The software was run in multimer mode using FASTA files as input(58, 59). Models were visualized and prepared for figures using PyMOL.

## Acknowledgements

M.A.B.B. acknowledges funding support from a Scientia Fellowship from UNSW, Human Frontiers Science Program Project Grant No. RGY0072/21 and U.S. Navy Office of Naval Research Research Grant No. N62909-22-1-2051.

