## Supplementary Figures 1-9, Supplementary Table 2 for "Hybrid Exb/Mot stators require substitutions distant from the chimeric pore to power flagellar rotation"

**Supplementary Material**

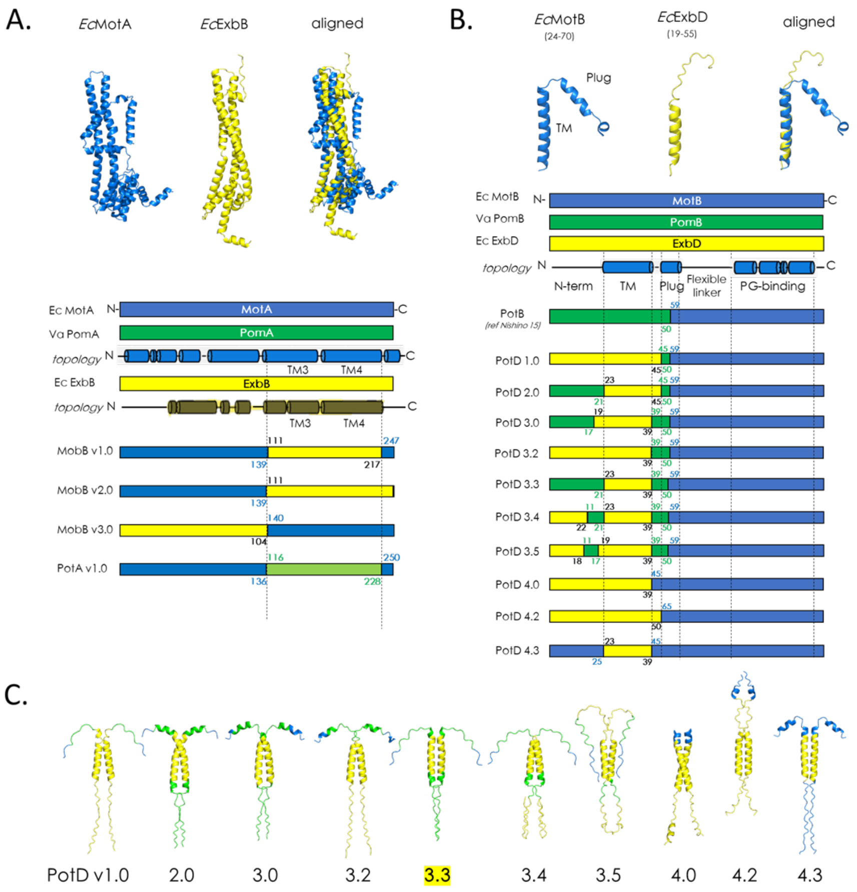

**Supplementary Figure 1. Design of Chimeric Exb/Mot complexes.**

Chimeric constructs of MotA/PomA/ExbB (A) and MotB/PotB/ExbD (B) were designed based on the topology of each subunit. The constructs shown in (A) were spliced at the junction between the N-terminal section of the protein and the more conserved TM3/4 domains. Similarly, the constructs in (B) were spliced at various points within the *N-terminal*, *TM* and *Plug* domains of the protein while retaining the *flexible linker* and *PG-binding* domains of MotA unchanged, based on a previously developed MotB/PomB chimera (PotB, Nishino et al. 2015). The amino acid residues at the junction points of each construct are indicated on each bar diagram and color-coded according to their identity (blue: MotA and MotB, green: PomA and PomB, yellow: ExbB and ExbD). C) Alphafold2 dimeric models of B-subunit chimeras (PotD variants) shown in **B** (with only *N-terminal*, *TM* and *Plug* domains shown), coloured according to their original. Construct PotD 3.3, which rescued the motility phenotype, is highlighted in yellow.

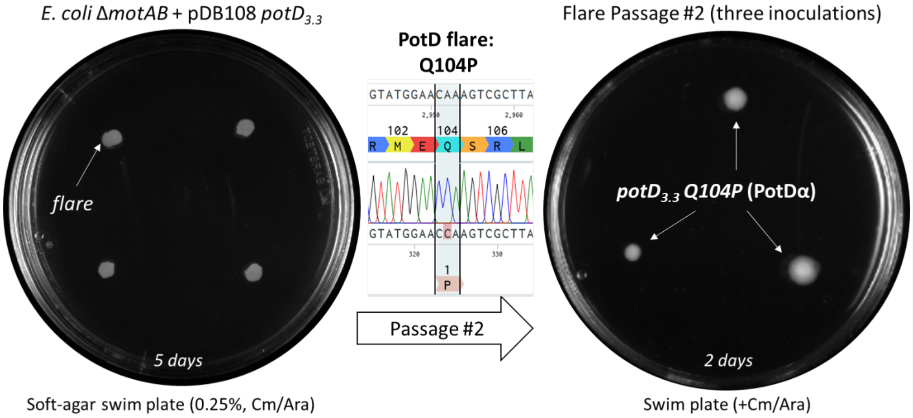

**Supplementary Figure 2**. **Adaptation of a motile subpopulation carrying a chimeric construct.** (left) Stator-deleted Δ*motAB E. coli* expressing PomA/PotD$_{3.3}$ were inoculated on a swim plate for five days. (centre) Extraction and re-sequencing of plasmid carrying the chimeric construct revealed a novel mutation (Q104P) in the *potD*$_{3.3}$ construct. The chromatogram output from Sanger sequencing highlights the codon C**A**A (Glutamine 104) in the original PotD$_{3.3}$ construct having mutated to C**C**A (Proline). (right) Triplicate inoculation of the mutant flare after two days in culture.

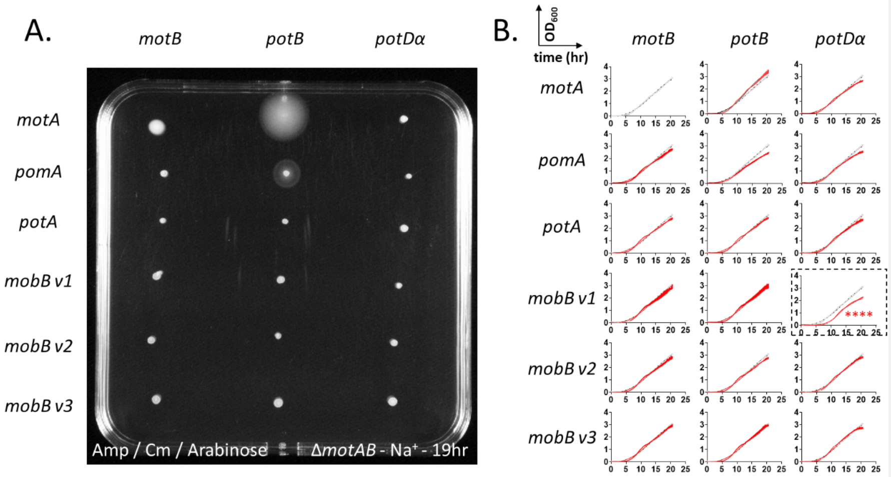

**Supplementary Figure 3. Swim-plate motility assay of *E. coli* co-expressing combinations of A and B subunits.**

(A) Soft agar swim plate (0.3% agar) (15 x 15 cm) inoculated with an array of stator-deleted *E. coli* Δ*motAB* clones co-expressing A- and B- subunits as indicated by their arrangement. Reagents and strains used in the assay on swim-agar (0.25%, 30 °C, 19 hr) are indicated at the bottom of the plate. Cells co-expressing MotA/MotB, MotA/PotB and PomA/PotB combinations showed a motile phenotype, while other combinations did not spread. (B) Growth profiles of strains shown in A, arranged in the same order, where x axis indicates time (hours) and y axis indicates optical density at 600nm ($OD_{600}$). Curves show the standard deviation at each time point from an average of three measurements. Each subunit combination is shown in red and overlayed onto the profile of the control strain *E. coli* Δ*motAB* expressing MotA/MotB (grey curve). Statistical significance was assessed using Student's *t*-test (**** $p < 0.0001$).

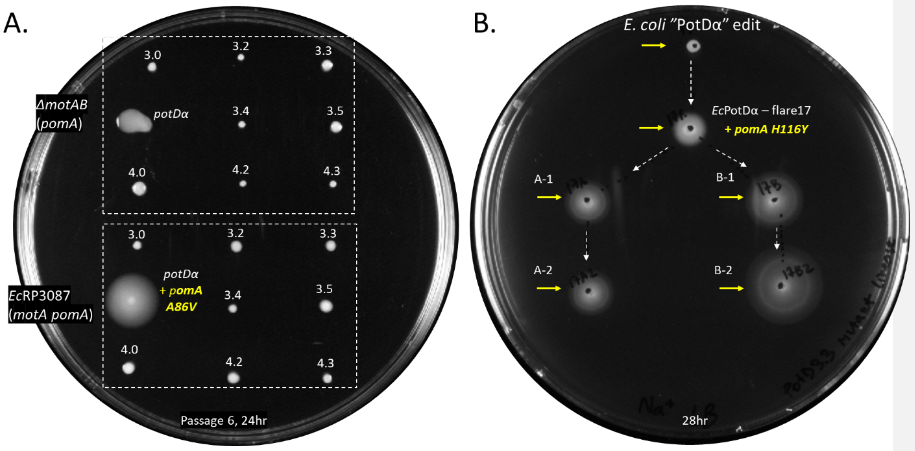

**Supplementary Figure 4. Optimization of motility via directed evolution of stator chimeras.**

(A) Motile phenotype of Δ*motAB* (top) and RP3087 (Δ*motB,* bottom) *E. coli* strains expressing B-subunit chimeras after 24 hr of culture on soft-agar plates (10 cm diameter). Clones were propagated on swim agar for 6 passages prior to the experiment shown here. Yellow labels indicate motile variants and any mutation found in their construct-bearing plasmids after resequencing. The upmotile *pomA* A86V mutation found in Δ*motB E. coli* RP3087 carrying PomAPotDα on plasmid is highlighted. (B) Motile phenotype of a synthetic lineage edited to replace the native *motA motB* locus of *E. coli* RP437 with *pomApotD*$_{3.3}$ Q104P (*E. coli* "PotDα") after 28 hr of culture on 0.3% soft agar. The original edited strain is shown at the top of the plate. Clones descending from repeated passages of motile subpopulations of the edited strain are shown below to recapitulate lineages A and B, linked by dotted arrows. Whole-genome sequencing of all six lineage members revealed the presence of the *pomA H116Y* mutation in all descendants of the original edit. A complete list of mutations found from WGS of all lineage members is reported in Supplementary Table 2.

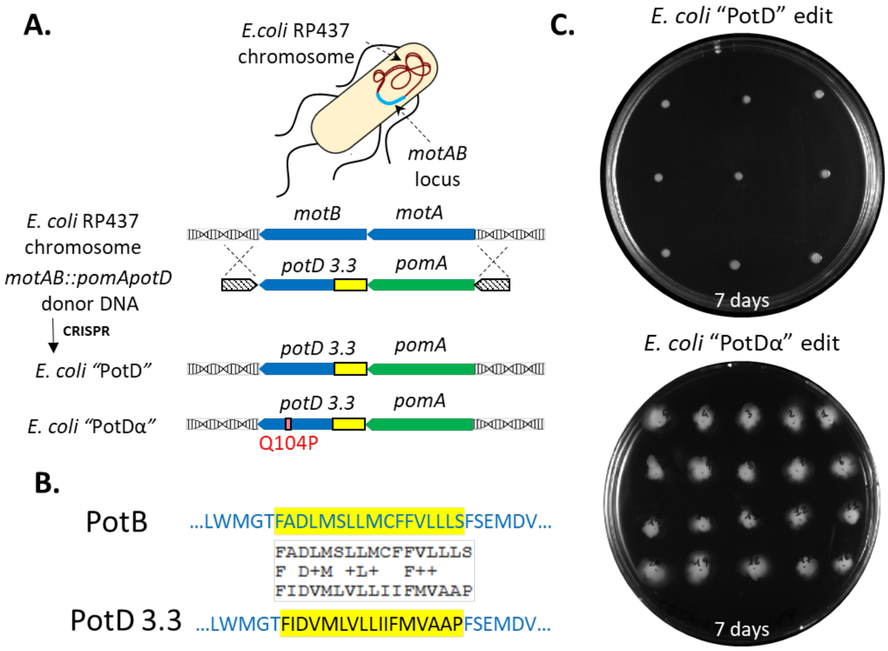

**Supplementary Figure 5. No-SCAR editing of *pomA* / *potD*$_{3.3}$ and its Q104P variant.**

(A) Schematic diagram of No-SCAR editing at the *motAB* locus on the *E. coli* RP437 chromosome to generate *E. coli* strains harbouring designed ("PotD") and evolved ("PotDα") PotD variants. (B) Highlight of the different amino acid sequences expressed in the template construct PotB and the PotD$_{3.3}$ construct spliced with ExbD (yellow highlight). Amino acid alignment highlights the degree of conservation between the two proteins. (C) Swim plate assays (7-day after inoculation) of multiple clones of No-Scar edited cells carrying the "PotD" (top) or "PotDα" (bottom) edits at the *motAB* locus.

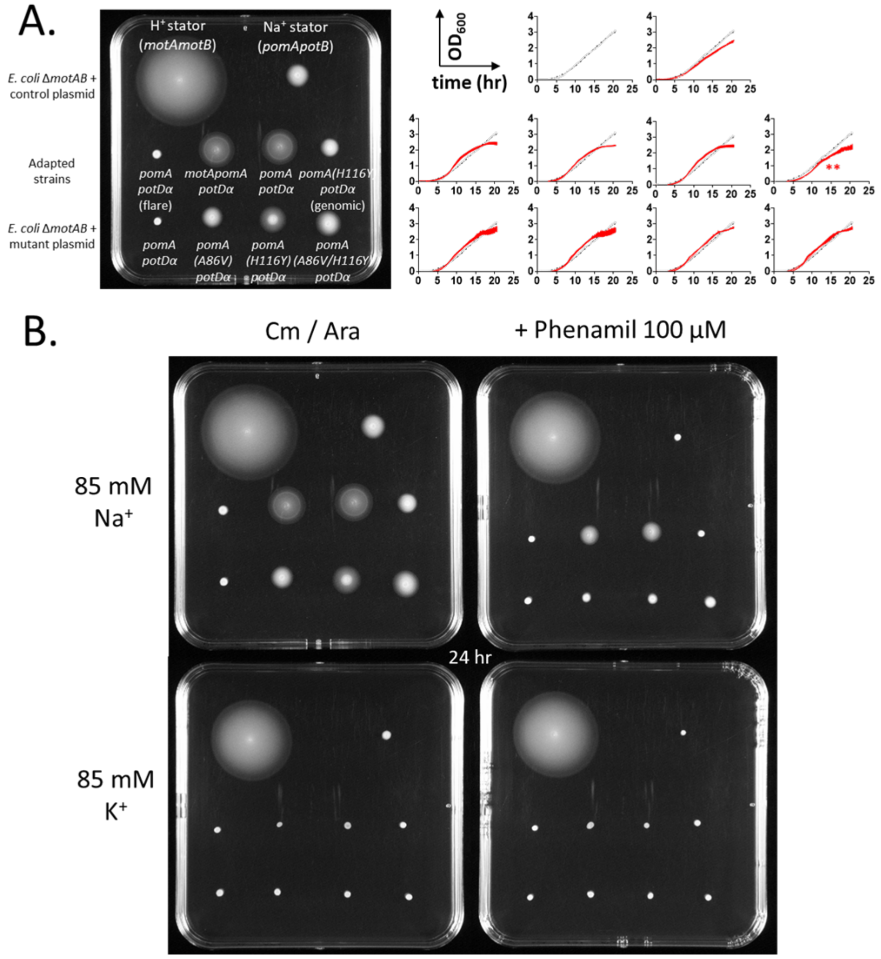

**Supplementary Figure 6. Phenotypic comparisons between strains obtained during the study.**

A) Swim-plate assay of strains collected over the course of the study, labelled above each inoculated colony. Growth profiles for each of the strains on the swim plate, arranged in the same order, where x axis indicates time (hours) and y axis indicates optical density at 600nm ($OD_{600}$), and overlayed onto the profile of control strain ($H^+$ stator MotAB). Statistical comparison performed using Student's T test, ** $P < 0.01$. (B) Swim plate assay on different substrates and arranged in the same order of (A). Plates containing either 85 mM $Na^+$ or 85 mM $K^+$ were supplemented with motility inhibitors for sodium-powered stators (100 µM Phenamil). Chimeric constructs were not motile on $K^+$ substrates.

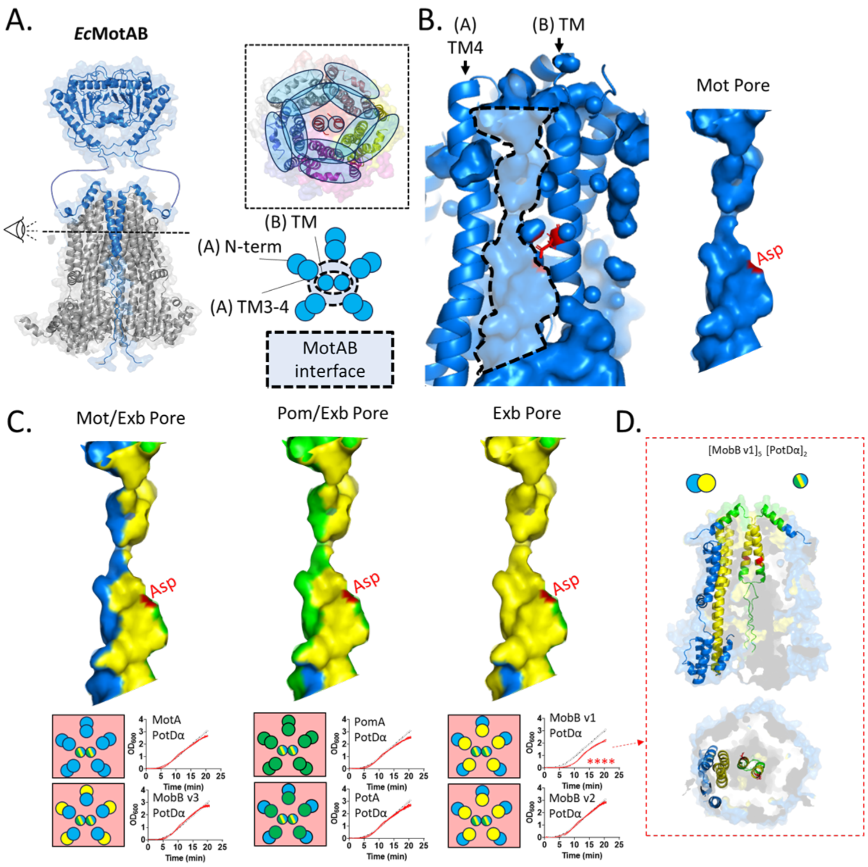

**Supplementary Figure 7. Visualization of chimeric pores and their impact on cell growth.**

(A) Side view and section of *Ec*MotAB (Alphafold2 model). The circle diagram highlights the expected locations of the ion-conducting pores at the subunit interface (blue donut with dashed black line). (B) Side view of the inter-subunit volume (pore) with catalytic aspartate side chain (Asp, D32) shown in red, displayed on the pore lumen. (C) Surface representation of ion-conduction pathways at the chimeric subunit interface. Subunit components are color-coded as in Fig. 2. Details from Fig. 3 and Supplementary Fig. 3 are combined below each pore surface model to show their architecture and effect on cell growth. A larger scale protein model is shown for the cytoplasmic and TM portion (PGB domain excluded here) of chimeric complex $[\text{MobB v1}]_5[\text{PotD}_{3.3}\ \text{Q104P}]_2$ shown to delay cell growth.

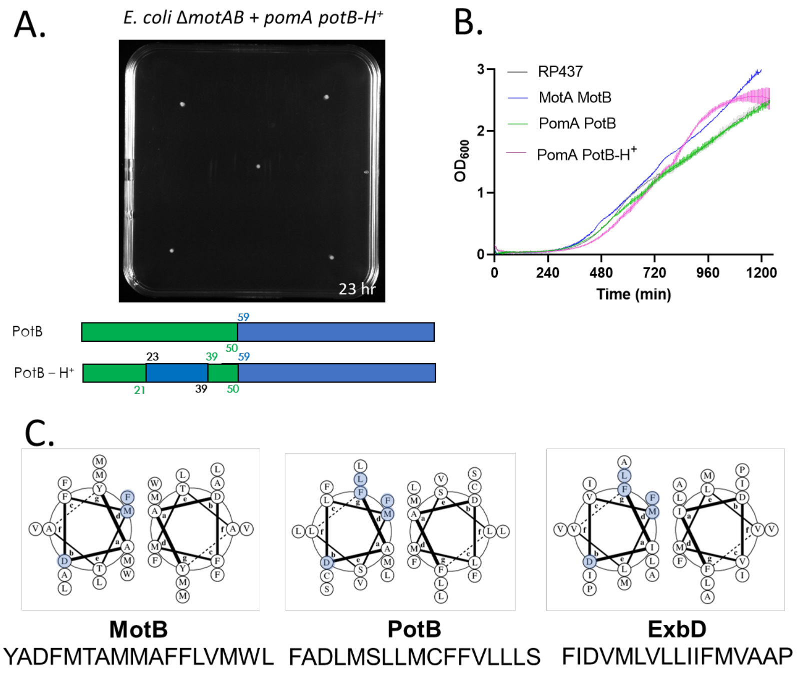

**Supplementary Figure 8. Testing a TM-swapped variant of PotB.** A) Soft agar motility assay for *E. coli* Δ*motAB* cells expressing *pomA* and *potB-H$^+$*. Five separate colonies were inoculated on soft agar but none developed swim rings after 23 hr of incubation. The combination of A and B stator subunit was deemed not functional. A schematic diagram of the PotB and the TM-swapped PotB-H+ constructs is shown underneath, coloured and arranged in the scheme of Fig. 2A. B) Overlaid growth curve profiles (Optical Density at 600 nm) for parent *E. coli* RP437 and *E. coli* Δ*motAB* expressing *motA motB* (blue), *pomA potB* (green) or *pomA potB-H$^+$*. C) Coiled-coil models (register *g*) of the TM domains of MotB, PotB and ExbD. Conserved residues are highlighted in blue on one of the two coils.

**Commented [PR1]:** •**Question #5. If ExbBD is actually an H+ (H3O+)-driven motor, it seems that a reasonable control would be to replace those same 17 residues of PotB with the corresponding residues from MotB. What would the conclusion be if that did or did not produce a functional chimera?** *The suggested chimeric design would be expected to support H+ conduction across the stator thus allowing for motility in low sodium (if able to couple with the BFM rotor). Proton leakage through an unplugged, but uncoupled, stator would be expected to slow the growth rate of the host due to dissipation of the PMF.* ***Our results show that the suggested TM-swapped B subunit does not allow for motility in a K+-based swim plate does not affect the growth rate of the E. coli host when compared to WT stators or control chimeras (when tested with PomA).***

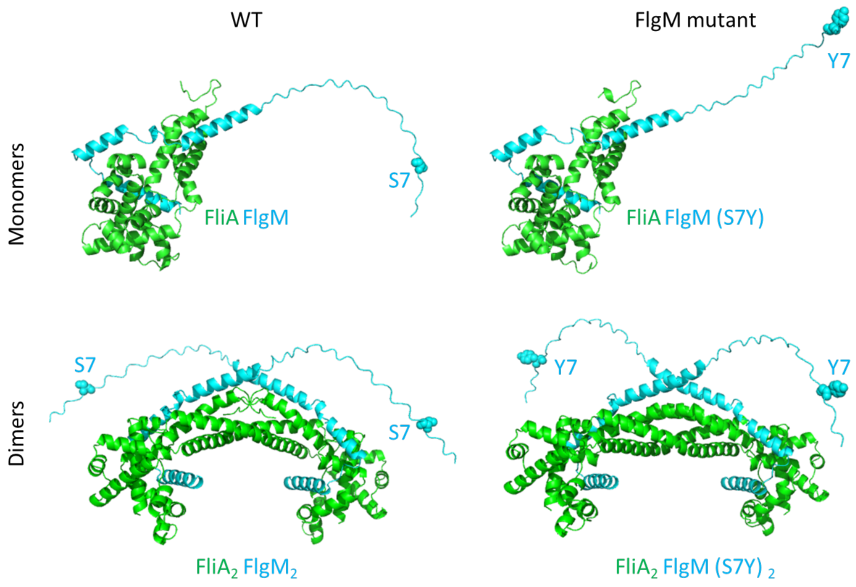

**Supplementary Figure 9. Alphafold2 models of *Ec* FliA FlgM and mutants.** Model structures of interacting *Ec* FliA (green) and *Ec* FlgM (cyan) monomers (top row) and dimers (bottom row). Wild-type proteins (left) or models including the FlgM S7Y mutant (right) are shown and the position of residue 7 of FlgM is highlighted by cyan spheres.

Commented [PR2]: •**Question #5. If ExbBD is actually an H+ (H3O+)-driven motor, it seems that a reasonable control would be to replace those same 17 residues of PotB with the corresponding residues from MotB. What would the conclusion be if that did or did not produce a functional chimera?** *The suggested chimeric design would be expected to support H+ conduction across the stator thus allowing for motility in low sodium (if able to couple with the BFM rotor). Proton leakage through an unplugged, but uncoupled, stator would be expected to slow the growth rate of the host due to dissipation of the PMF.* ***Our results show that the suggested TM-swapped B subunit does not allow for motility in a K+-based swim plate does not affect the growth rate of the E. coli host when compared to WT stators or control chimeras (when tested with PomA).***

**Supplementary Table 1. Oligonucleotide and gene fragment sequences.**

List of primers used in No-SCAR editing of *E. coli* RP437 to Knock-in chimeric constructs or knock-out the native *motAB* locus. List of primers used to introduce mutations A86V and H116Y in *potD*$_{3.3}$ encoded on a pSHU1234 plasmid using the Quikchange® method. List of synthetic gene fragments (gBlocks) encoding the chimeric constructs described in this study. List of primers used in the cloning of the PotB-H+ chimera via inverse PCR.

| **E.coli "PotDα" edit** | | | | | | | |
|---|---|---|---|---|---|---|---|
| REFERENCE CHROMOSOME | POS | TYPE | REF | ALT | EVIDENCE | EFFECT | GENE |
| CP083410.1b (E.coli "PotDα" edit) | none | | | | | | |
| **E.coli "PotDα" edit - flare17** | | | | | | | |
| REFERENCE CHROMOSOME | POS | TYPE | REF | ALT | EVIDENCE | EFFECT | GENE |
| CP083410.1b (E.coli "PotDα" edit) | 1130109 | snp | G | T | T:33 G:0 | S7Y | flgM |
| CP083410.1b (E.coli "PotDα" edit) | 1196277 | complex | CGCGAAA | TGCCAAG | TGCCAAG:10 CGCGAAA:0 | | icd |
| CP083410.1b (E.coli "PotDα" edit) | 1976659 | snp | G | A | A:35 G:0 | H116Y | pomA |
| **E.coli "PotDα"_A-1** | | | | | | | |
| REFERENCE CHROMOSOME | POS | TYPE | REF | ALT | EVIDENCE | EFFECT | GENE |
| CP083410.1b (E.coli "PotDα" edit) | 1130109 | snp | G | T | T:38 G:0 | S7Y | flgM |
| CP083410.1b (E.coli "PotDα" edit) | 1976659 | snp | G | A | A:44 G:0 | H116Y | pomA |
| **E.coli "PotDα"_A-2** | | | | | | | |
| REFERENCE CHROMOSOME | POS | TYPE | REF | ALT | EVIDENCE | EFFECT | GENE |
| CP083410.1b (E.coli "PotDα" edit) | 1130109 | snp | G | T | T:28 G:0 | S7Y | flgM |
| CP083410.1b (E.coli "PotDα" edit) | 1976659 | snp | G | A | A:31 G:0 | H116Y | pomA |
| **E.coli "PotDα"_B-1** | | | | | | | |
| REFERENCE CHROMOSOME | POS | TYPE | REF | ALT | EVIDENCE | EFFECT | GENE |
| CP083410.1b (E.coli "PotDα" edit) | 1130109 | snp | G | T | T:38 G:0 | S7Y | flgM |
| CP083410.1b (E.coli "PotDα" edit) | 1976659 | snp | G | A | A:43 G:0 | H116Y | pomA |
| **E.coli "PotDα"_B-2** | | | | | | | |
| REFERENCE CHROMOSOME | POS | TYPE | REF | ALT | EVIDENCE | EFFECT | GENE |
| CP083410.1b (E.coli "PotDα" edit) | 1130109 | snp | G | T | T:34 G:0 | S7Y | flgM |
| CP083410.1b (E.coli "PotDα" edit) | 1976659 | snp | G | A | A:31 G:0 | H116Y | pomA |

**Supplementary Table 2. Whole genome sequencing of lineage members of the *E. coli* "PotDα" family presented in Figure 4.**

The table shows the output of single nucleotide polymorphism analysis performed using Snippy (see Methods) using the whole-genome sequencing data obtained from clones shown in Supplementary Figure 4B (yellow arrows). Mutations were mapped on reference chromosome CP083410.1b, a derivative of *CP083410*.1 (Ridone et al, 2022) encoding the *potD*$_{3.3}$ *Q104P* (PotDα) and *pomA* at the *pomA/potB* locus (originally replacing the native *motAB* locus *of E. coli* RP437).
